# Cryo-EM structure of the human Sirtuin 6-nucleosome complex

**DOI:** 10.1101/2023.03.17.533206

**Authors:** Un Seng Chio, Othman Rechiche, Alysia R. Bryll, Jiang Zhu, Jessica L. Feldman, Craig L. Peterson, Song Tan, Jean-Paul Armache

## Abstract

Sirtuin 6 (SIRT6) is a multifaceted protein deacetylase/deacylase and a major target for small-molecule modulators of longevity and cancer. In the context of chromatin, SIRT6 removes acetyl groups from histone H3 in nucleosomes, but the molecular basis for its nucleosomal substrate preference is unknown. Our cryo-electron microscopy structure of human SIRT6 in complex with the nucleosome shows that the catalytic domain of SIRT6 pries DNA from the nucleosomal entry-exit site and exposes the histone H3 N-terminal helix, while the SIRT6 zinc-binding domain binds to the histone acidic patch using an arginine anchor. In addition, SIRT6 forms an inhibitory interaction with the C-terminal tail of histone H2A. The structure provides insights into how SIRT6 can deacetylate both H3 K9 and H3 K56.

**Teaser:** The structure of the SIRT6 deacetylase/nucleosome complex suggests how the enzyme acts on both histone H3 K9 and K56 residues.

## Introduction

Sirtuins are evolutionarily conserved metabolic sensor enzymes that use NAD+ as a coenzyme (*1, 2*). A critical role of sirtuins in aging was first suggested by studies of yeast Sir2 (*3*). In mammals, depletion of the SIRT6 sirtuin results in shortened lifespans while SIRT6 overexpression extended lifespans (*4–7*). Additionally, SIRT6 is associated both with tumor suppression and tumorigenesis in different cancers (*8*). These aging and cancer-related phenotypes are linked to the ability of SIRT6 to deacylate substrates such as tumor necrosis factor-alpha (TNF-α) (*9*) and to deacetylate histone H3K9ac and H3K56ac on nucleosomes (*10,11*). Histone H3K9ac is a mark of transcriptionally active promoters (*12*) and histone H3K9 acetylation plays an important role in DNA repair and telomere maintenance. Crystal structures and biochemical studies have explored how SIRT6 binds a peptide substrate, its NAD^+^ cofactor and allosteric effectors (*9, 13–20*). However, the molecular basis for the sequence preference of SIRT6 to deacetylate histone H3 at positions K9 and K56 (*10, 21, 22*) is not known. Nor do we understand the structural basis for the preference of SIRT6 to deacetylate nucleosomes at H3 K9 and K56 over free histones (*23*). To address these deficiencies, we have determined the structure of SIRT6 in complex with its nucleosome substrate. The multivalent interactions between SIRT6 and both histone and DNA components of the nucleosome provide insights into how SIRT6 can deacetylate exposed H3 K9 and occluded H3 K56 residues.

## Results

### SIRT6-nucleosome structure determination

We obtained 2.7 – 3.1 Å resolution cryo-EM maps of SIRT6 in complex with a 172 bp nucleosome containing a 26 bp DNA extension on one end **(Fig. 1, Figs. S1 & S2)**. Although our sample was prepared in the presence of SNF2h, which SIRT6 recruits to sites of DNA damage (*24*), we do not see density for SNF2h. This suggests that the ATPase may engage with either the nucleosome and/or SIRT6 in a transient or flexible manner. We also do not see density for the intrinsically disordered C-terminal domain of SIRT6, which was previously reported to bind nucleosomal DNA (*25*). This does not exclude the possibility that dynamic interactions between the SIRT6 C-terminal domain and nucleosomal DNA were not resolved by single particle cryo-EM. Our data also yielded a subset of particles containing two copies of SIRT6 simultaneously bound to opposite faces of the nucleosome **(Fig. S2)**. The density for the SIRT6 bound to the face associated with the DNA extension appears much stronger relative to the density for the SIRT6 on the opposite face without extended DNA, suggesting that the presence of extended DNA stabilizes the positioning of SIRT6 on the nucleosome.

**Fig. 1:**
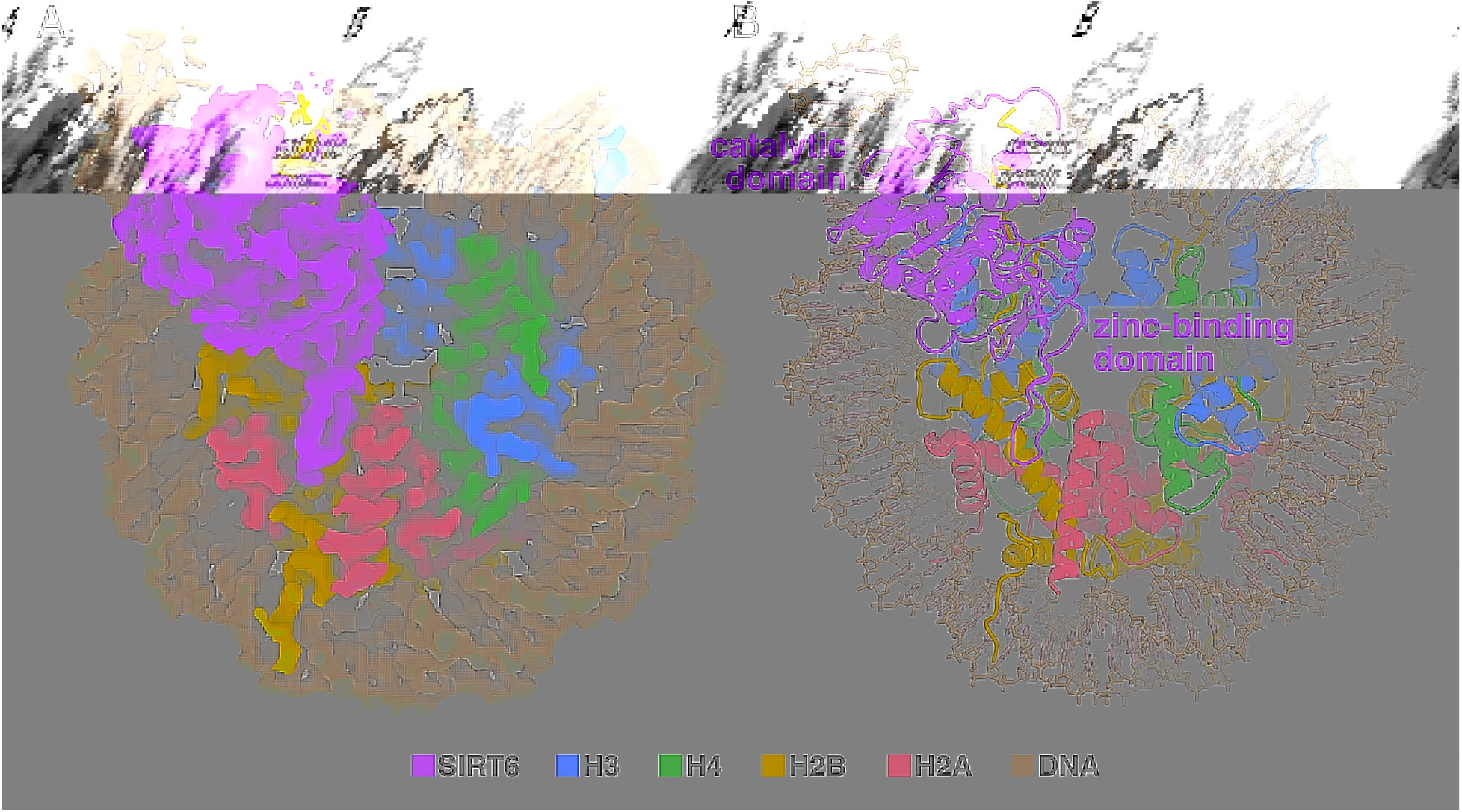
Overview of SIRT6/nucleosome structure. (**A**) 3.07 Å cryo-EM Coulomb potential density map of structure, (**B**) Cartoon representation of structure.

### Overview of SIRT6-nucleosome complex

Our study shows that the SIRT6 deacetylase domain forms multivalent interactions with the nucleosome via the nucleosome acidic patch, the H3 N-terminal histone tail, the C-terminal H2A tail and nucleosomal DNA **(Fig. 1)**. The structure of SIRT6 is very similar in the absence or presence of the nucleosome except for residues in and around the SIRT6 NAD^+^ binding loop and separately, the 10 residues that form the SIRT6-specific extended loop in its zinc-binding domain (*13*). The orientation of the SIRT6 zinc-binding motif with respect to its catalytic Rossman-fold domain is unchanged upon nucleosome binding. In contrast, the nucleosome is partially unwrapped, with DNA displaced from the extended end of the nucleosome to accommodate SIRT6-DNA interactions.

### SIRT6-nucleosome acidic-patch interaction

Previous biochemical experiments indicate that SIRT6 engages with a nucleosome through the acidic patch formed by histones H2A and H2B (*25*), and hydrogen-deuterium exchange mass spectrometry experiments suggested that an N-terminal helix of SIRT6 (residues 28-43) may participate in nucleosome interactions (*25*). However, our structure shows that SIRT6 binds to the acidic patch using the extended loop (^167^TVAKARGLRA^176^) **(Fig. 2A)** in its zinc-binding domain that is absent in other sirtuins (*13*). In the previous hydrogen-deuterium exchange experiments (*25*), peptides for zinc-binding domain were not observed, which likely prevented assessing whether this region interacts with the nucleosome. The zinc-binding domain can be thought of as a pipe that fills the shallow ditch formed by the H2A/H2B acidic patch. These contacts are mediated by a combination of hydrophilic and van der Waals interactions. Prominent among these interactions is SIRT6 R175 which makes an arginine anchor interaction with H2A acidic patch residues E61, D90 and E92. In addition, other SIRT6 basic residues interact with the acidic patch. SIRT6 R172 constitutes a type 1 variant arginine and interacts with H2A E56, H2B Q44 and E110, while SIRT6 R178 acts as an atypical arginine to interact with H2A D90 and E92 following the arginine-acidic patch nomenclature proposed by McGinty and Tan (*26*). In addition, SIRT6 K170 makes ionic interactions with H2A E64. These contacts are consistent with biochemical data where mutation of H2A residues E61, E64, D90, and E92 to alanines resulted in weaker SIRT6 binding (*25*). Mutating each of these SIRT6 basic side chains (K170, R172, R175 and R178) to alanine reduced binding to nucleosomes **(Fig. 2B)**, with R175 showing the largest effect with ~9-fold weaker binding consistent with its critical role as an arginine anchor. Furthermore, the SIRT6(R175A) has little or no H3K9ac nucleosomal deacetylation activity, corroborating the importance of the R175 arginine anchor for SIRT6 enzymatic activity **(Fig. 2C)**.

**Fig. 2:**
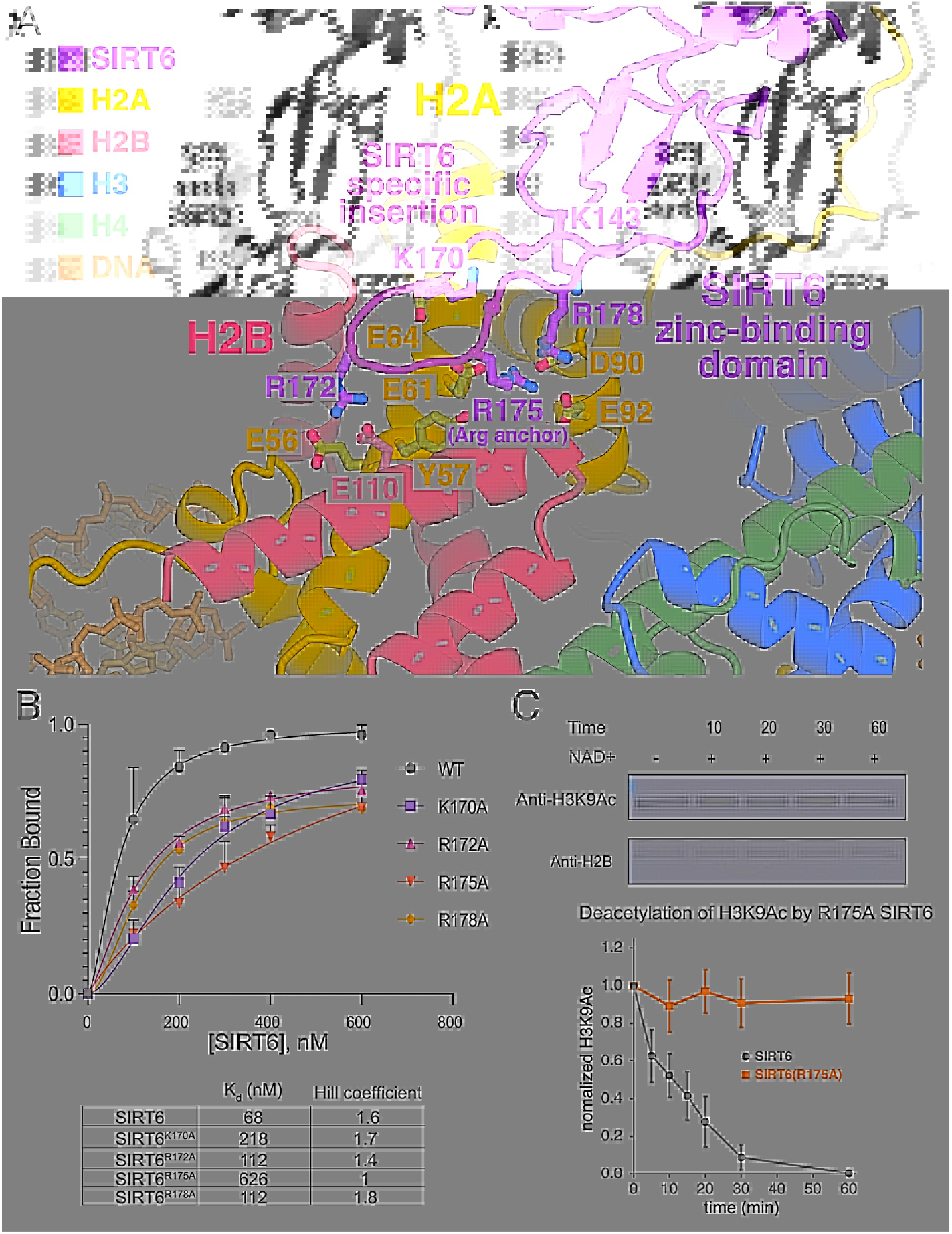
Interactions of SIRT6 zinc-binding domain with nucleosome acidic patch. (**A**) Cartoon representation of SIRT6 zinc-binding domain with histone dimer acidic patch, (**B**) Quantification of nucleosome binding assay for wild type and mutant SIRT6, (**C**) SIRT6 H3K9Ac deacetylase assay for wild type and R175A SIRT6 normalized against H2B with representative Western blot (top) and plot showing standard deviation error bars with n = 3 (bottom).

### SIRT6-H2A tail interaction

In most nucleosome complex structures, the H2A C-terminal tail is disordered beyond K118 or K119. In our SIRT6-nucleosome structure, weak density for approximately 10 more residues of the histone H2A C-terminal tail is visible and tracks upwards to interact with the SIRT6 catalytic domain **(Fig. 3A)**. This interaction was not predicted in previous SIRT6 studies and to the best of our knowledge, this is the first example of a chromatin enzyme interacting with the H2A C-terminal tail. To test the importance of the H2A C-terminal tail interaction, we reconstituted nucleosomes with histone H2A lacking the C-terminal tail and tested the ability of SIRT6 to deacetylate H3K9ac on these nucleosomes. Intriguingly, we observe increased SIRT6 H3K9ac deacetylation activity on nucleosomes lacking the H2A C-terminal tail **(Fig. 3B)**. This suggests that the H2A tail may have an inhibitory role in regulating SIRT6 deacetylation activity on the nucleosome.

**Fig. 3:**
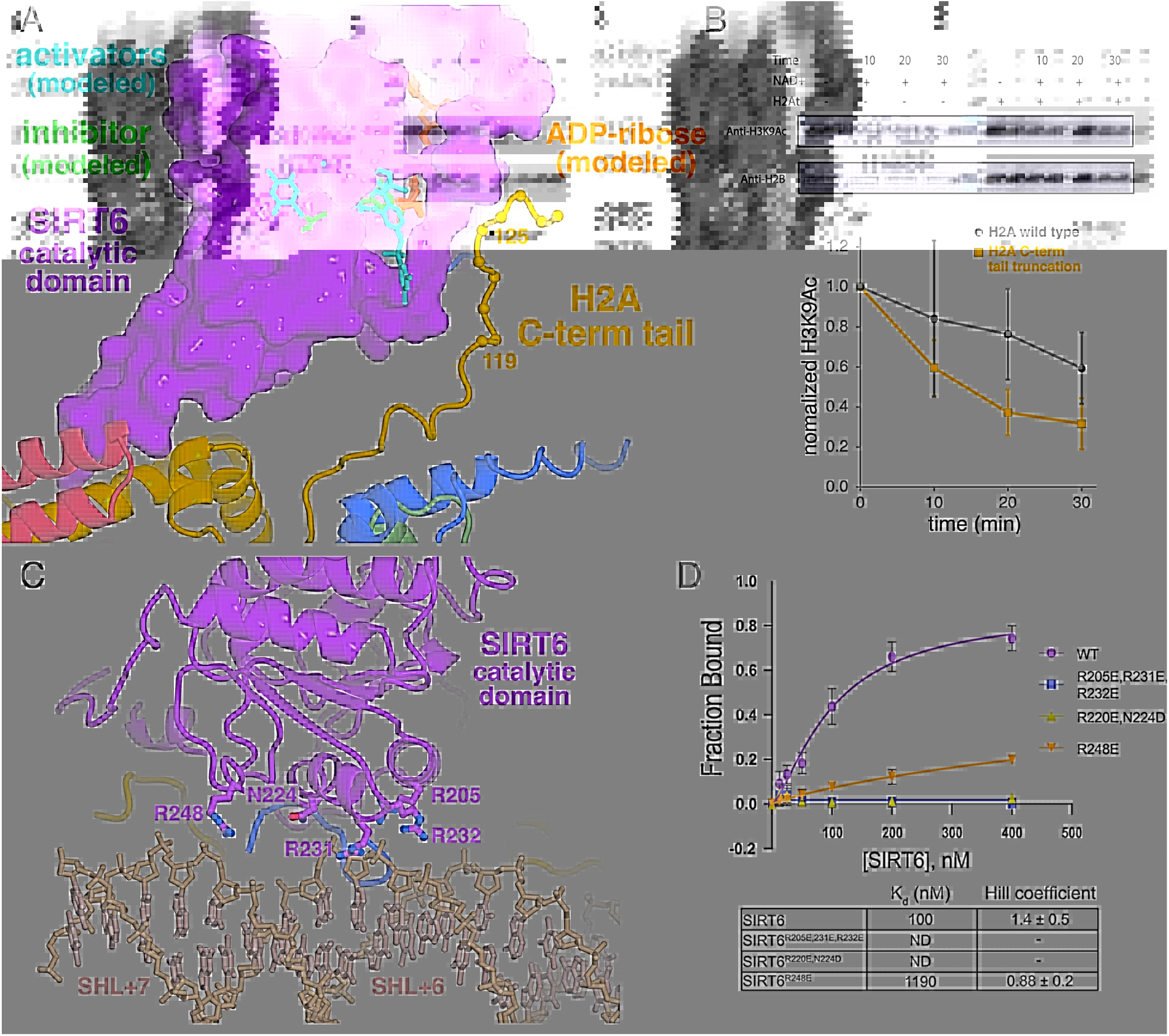
Inhibitory interactions of histone H2A C-terminal tail near SIRT6 allosteric binding pocket, and nucleosomal DNA interactions of SIRT6 catalytic domain with nucleosomal DNA. (**A**) The histone H2A C-terminal binds to SIRT6 proximal to allosteric activators (MDL-801, modeled from two different structures (PDB 5Y2F and 6XV1) and an allosteric inhibitor (catechin gallate, PDB 6QCJ). The modeled product analog, 2’-*O*-acyl-ADP-ribose, adopts a well-defined conformation in these three SIRT6-allosteric effector structures. The C_α_ positions of the H2A C-terminal residues 119-128 are shown as yellow spheres. (**B**) Deletion of the H2A C-terminal tail enhances SIRT6 nucleosomal H3K9Ac deacetylase activity (normalized against histone H2B). Representative Western blot data (top) and plot for HDAC assays (n=3) shown. (**C**) Interactions of SIRT6 catalytic domain with nucleosomal DNA. Same color code as for Fig. 2a. (**D**) Quantification of nucleosomal binding assay for wild type and SIRT6 mutated in DNA-binding residues

Our SIRT6-nucleosome sample used for structural determination lacks NAD^+^, which is a necessary co-factor for SIRT6 deacetylation activity (*27*). In comparison to crystal structures of SIRT6 bound to ADP-ribose (*9, 13–20*), we notice that the density for SIRT6 residues 64-80 is missing in our map, although they are resolved in the crystal structures without nucleosomes **(Fig. 3A, Fig. S3)**. This region of SIRT6 contains residue R65, which makes extensive contacts with the adenosine ribose and pyrophosphate of ADP-ribose in solved crystal structures and is important for the activation of SIRT6 for catalysis (*28*).The lack of defined density in our map suggests that this region is flexible in the absence of a bound co-factor and only becomes stabilized after co-factor binding. Previous biochemical data showing that R65 mediates a necessary conformational change for activation supports this interpretation (*28*).

### SIRT6-nucleosome DNA interactions

A previous prediction suggested that the disordered SIRT6 CTD can bind DNA (*25*). We find instead that the SIRT6 catalytic domain interacts with nucleosomal DNA. We observe multiple arginine residues within the SIRT6 catalytic domain (R205, R220, R231, R232, R248) that contact nucleosomal DNA near the entry-exit site of the nucleosome. R205, R231 and R232 contact the DNA phosphate backbone at superhelical location 6 (SHL+6) while R248 binds across the major groove to the opposite DNA strand phosphate backbone at SHL+7 just beyond the nucleosome core **(Fig. 3C)**. The polar residue, N224, in the vicinity may also have a role in the interaction. These interactions are made only to the DNA phosphate backbone with no apparent contacts to the nucleotide bases. To determine the contribution of these residues to SIRT6-nucleosome binding, we generated three sets of SIRT6 mutants: an R205E/R231E/R232E triple mutant, an R220E/N224D double mutant, and an R248E point mutant. SIRT6(R248E) binding to nucleosomes was severely impaired (~12-fold weaker vs. wild-type), and the SIRT6 triple and double mutants were no longer able to bind nucleosomes at all **(Fig. 3D)**. These results suggest that the SIRT6 globular domain interactions with nucleosomal DNA play a critical role in stabilizing the SIRT6-nucleosome complex. We also generated and visualized two new SIRT6-nucleosome complexes, containing shorter DNA constructs (147 and 145 bp, **Figs. S4 & S5**). These reconstructions revealed significantly reduced quality of SIRT6, thereby further suggesting that DNA-SIRT6 interactions play a significant role in the stabilization of the complex.

### Positioning of H3 substrate residues

We have built H3 residues 3-12 into relatively weak density occupying the same peptide substrate binding site in previous SIRT6/myristoylated H3 peptide crystal structures (*9, 15*). In our structural model, the conformation of the H3 tail substrate is similar but not identical to the SIRT6/myristoylated H3 peptide structures with the H3 K9 side chain positioned essentially equivalent to the myristoylated H3 K9. The H3 tail substrate is sandwiched between the SIRT6 active site and nucleosomal DNA, allowing the H3 K4 side chain to fill the DNA minor groove at SHL 6.5 **(Fig. 4A)**. We wondered if this H3 K4-DNA interaction and potential interactions made by H3 residues between K4 and K9 might account for the H3K9 histone deacetylase specificity of SIRT6. We therefore assayed the histone deacetylase activity of SIRT6 on H3 K9 acetylated nucleosomes also containing H3 mutated at specific tail residues. Mutating the H3 K4 to glutamic acid slightly decreased the ability of SIRT6 to deacetylate H3 K9 **(Fig. 4B)** in nucleosomes. Similar modest adverse effects on SIRT6 H3 K9 deacetylation were observed when the H3 R8 side chain, which might interact with SIRT6, was removed and when one residue between H3 K4 and K9, Q6, was deleted. These results suggest a role of H3 residues 4-8 in SIRT6 deacetylation of H3 K9, but further investigation will be needed to determine what defines the specificity of SIRT6 for H3 K9.

**Fig. 4:**
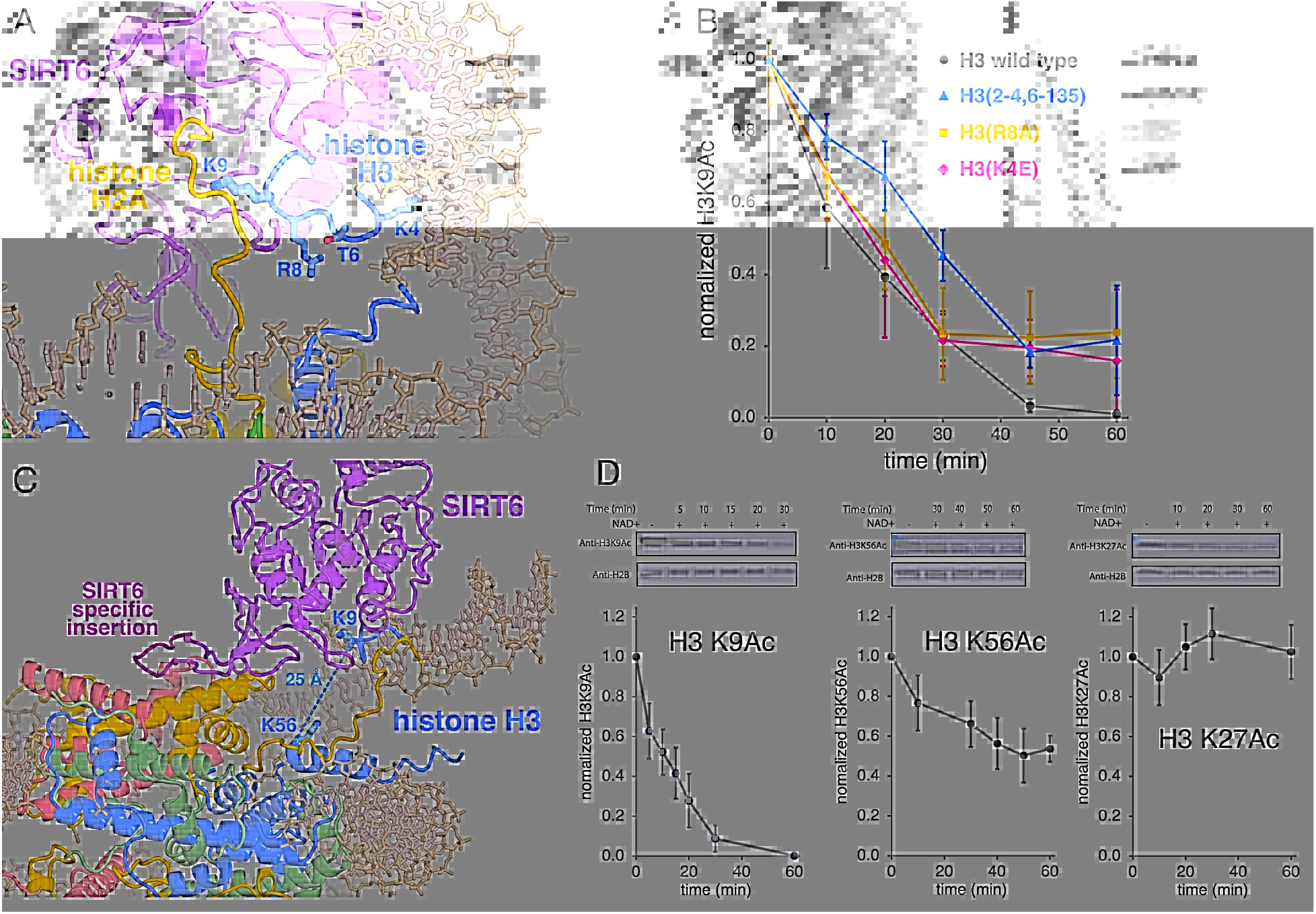
SIRT6 nucleosomal substrate specificity. (**A**) Cartoon and stick representation of SIRT6 binding of H3 tail around substrate residue K9. (**B**) Point mutations and deletions of H3 tail residues that interact SIRT6 adversely affect SIRT6 nucleosomal histone H3K9Ac deacetylase activity. (**C**) Cartoon and stick representation of SIRT6/nucleosome complex shows that histone H3 K56 is exposed but is at least 25 Å from H3 K9 and the SIRT6 catalytic site. (**D**) SIRT6 HDAC activity on nucleosomal histone H3 K9, K56 and K27.

Although we observe H3K9 in the SIRT6 active site, H3K56 is far from SIRT6 active site **(Fig. 4C)**. In agreement with previous biochemical results (*23*), we observe that SIRT6 deacetylates H3K56ac albeit less efficiently compared to H3K9ac **(Fig. 4D, middle vs left)**. Our structural observation of H3K56 being distant from the SIRT6 active site raises the question of how SIRT6 accesses H3K56 for deacetylation (discussed further below). We further note that while a previous study proposed that nucleosomal H3K27ac is also a substrate for SIRT6 (*29*), we find that SIRT6 does not deacetylate nucleosomal H3K27ac **(Fig. 4D, right)**.

## Discussion

We have determined the cryo-EM structure of the SIRT6 histone deacetylase in complex with the nucleosome. Our structure explains how SIRT6 deacetylates histone H3 K9 in its physiological nucleosome substrate more efficiently than in peptide substrates. We observe that SIRT6 binds at the nucleosome entry-exit site, using its globular domain to pry DNA from the histone octamer. This action is facilitated by SIRT6’s multivalent interactions with both histone and DNA components of the nucleosome using its catalytic and zinc-binding domains which are rigidly attached to each other. On one end, SIRT6 uses its zinc-binding domain to dock onto the histone acidic patch. On the other end, SIRT6’s catalytic domain binds to nucleosomal DNA at the entry-exit site replacing contacts otherwise made by the histone H3 N-terminal helix and thus partially exposing this helix. This release also allows the histone H3 N-terminal tail region to position the target H3 K9 side chain into the SIRT6 catalytic site while simultaneously inserting the H3 K4 side chain into the nucleosome DNA minor groove at SHL 6.5. Thus, SIRT6’s multiple interactions with the nucleosome likely facilitates productive binding of the H3 tail for catalysis.

The histone dimer acidic patch is a frequent target of chromatin enzymes and factors with an arginine anchor often used to bind this nucleosomal patch. SIRT6 uses R175 in the zinc-binding domain as an arginine anchor to bind to the histone dimer acidic patch, with the arginine side chain matching the tightly clustered conformation of arginine anchors in other chromatin complexes (*26*). The critical importance of the R175 arginine anchor for SIRT6 function is confirmed by the approximate 10-fold reduction binding affinity to nucleosomes and apparent complete loss of H3 K9 deacetylase activity. We also find that mutations of SIRT6 zinc-binding domain basic residues K170, R172 and R178 adversely affect nucleosome binding, consistent with their interactions with the histone dimer acidic patch observed in our cryo-EM structure.

Unlike H3 K9 which resides in the unstructured H3 N-terminal tail, H3 K56 is located on the H3 N-terminal helix which interacts with nucleosomal DNA in the absence of SIRT6. It was previously unclear how SIRT6 would then be able to access the H3 K56 side chain for catalysis. Our structure shows that SIRT6 prying apart nucleosomal DNA from the histone octamer also exposes the H3 N-terminal helix **(Fig. 4C)**. Conversely, acetylation of H3 K56 increases unwrapping of entry-exit nucleosomal DNA (*30, 31*). In our structure, the alpha carbons of H3K56 and H3K9 are 25 Å apart and it is clear that in this state H3 K56 cannot reach the SIRT6 active site. We propose that the conformational flexibility in the 10 residue insertion of the SIRT6 zinc-finger domain might allow SIRT6 to pivot as a rigid body about the histone acidic patch to approach the histone core. This and the possible unwinding of the H3 N-terminal helix might allow H3K56 to enter the SIRT6 catalytic site. The additional distortions necessary for this to occur could explain the lower deacetylase activity on H3K56 substrates.

The H2A C-terminal tail interacts with a SIRT6 short helix adjacent to both the SIRT6 NAD+ binding loop (residues 55-65) and the allosteric binding pocket targeted by both activators MDL-801 and quercetin activators and the catechin gallate inhibitor **(Fig. 3A)**. For this reason and our observation of the inhibitory effect of the SIRT6-H2A tail interaction, we suspect that this structure represents SIRT6 bound to a nucleosome in an inhibited, non-activated state. Since the unbound structure of SIRT6 is largely unchanged when activated or inhibited by the allosteric effectors, it is possible or even likely that activation of SIRT6 will involve local conformational changes near the catalytic site without major changes in how SIRT6 binds to the nucleosome via interactions with the histone acidic patch or nucleosomal DNA. This underscores the significance of the SIRT6 allosteric binding pocket as a target for drug discovery as well as the importance of using physiological nucleosome substrates for characterizing the effect of drug candidates on the enzymatic activity of SIRT6.

Our structural and biochemical studies explain how multivalent interactions of SIRT6 with both histone and DNA components of the nucleosome enable SIRT6 to deacetylate H3K9ac. It should be noted that the SIRT6-nucleosome used for our structural studies did not include NAD+ or a cofactor analog. We suspect this is not a serious limitation since the SIRT6 catalytic domain adopts the same structure in the presence or absence of cofactor (*13, 32*). In terms of substrate specificity, our structure suggested possible mechanistic roles for specific H3 residue side-chains N-terminal to the H3 K9 target residue. However, the modest effect of mutating these residues on the H3K9ac deacetylase activity of SIRT6 suggests either a concerted role for these N-terminal residues or a more complicated explanation for why SIRT6 targets H3 K9 versus other lysines present in the unstructured H3 tail. The relatively weak density for the H3 tail peptide also limits our confidence in the quality of the structure in this particular region. Our SIRT6-nucleosome atomic model provides the structural framework for further studies to understand H3K9 specificity and remaining issues such how SIRT6 engages H3K56 for deacetylation and the role of the H2A tail in SIRT6 function.

## Materials and Methods

### Nucleosome preparation

Three DNA constructs centered on the Widom 601 sequence (*33*) were used for nucleosome assembly; these constructs consisted of 145 (0-145-0), 147 (1-145-1), and 172 (1-145-26) base pairs of DNA. Recombinant H3 and H4 *Xenopus laevis* histones and H2A and H2B human histones were expressed, purified, and reconstituted with each DNA construct as described previously (*34*), including anion-exchange purification of the nucleosomes.

### Protein purification

The gene coding for the full-length human SIRT6 (UniProtKB: Q8N6T7) was cloned into pST50Tr (*35*) vector with an N-terminal GSS-hexahistidine (His_6_). GSS-(His_6_)-SIRT6 was expressed in BL21(DE3) pLysS *Escherichia coli* cells at 23°C. Bacterial cells were lysed by sonication, and the crude lysate centrifuged at 36,000 g for 40 min at 4°C. The protein was purified by metal-affinity chromatography (Talon resin, Clontech), the affinity tag removed using tobacco etch virus (TEV) protease and the protein further purified by Source S cation-exchange chromatography (Cytiva).

His6-tagged human SNF2h was expressed and purified as previously described with minor modifications (*36*). Briefly, His6-SNF2h was expressed in BL21(DE3) Rosetta *Escherichia coli* cells at 18°C. Cells were lysed via sonication, and Ni-NTA affinity chromatography used to isolate His6-SNF2h from the clarified lysate. TEV protease was used to remove the His6-tag, and the untagged SNF2h was passed through a HiTrapQ column (Cytiva) to remove contaminating DNA. The protein was then run over a HiLoad Superdex200 column (Cytiva), and pure fractions were pooled, aliquoted, and stored at −80 °C.

### Cryo-EM sample preparation

SIRT6 was reconstituted in reconstitution buffer (20 mM HEPES pH 7.5, 75 mM NaCl, 1 mM DTT) at 0.8:1 enzyme:nucleosome ratio with the 145 and 147 base pair nucleosomes, and at 2.2:1 ratio with the 172 base pair nucleosomes. SNF2h was also included in the SIRT6-172 base pair nucleosome sample at 1.1-fold excess over nucleosome.

The SIRT6-147 base pair nucleosome sample was crosslinked in reconstitution buffer with 0.05% glutaraldehyde. The sample was incubated on ice for 10 minutes and then quenched with 100 mM Tris-HCl pH 7.5. The SIRT6-145 base pair / SIRT6-172 base pair nucleosome samples were first purified using a Superdex 200 column (GE Healthcare) and then stabilized using the GraFix method (*37*). Light buffer (20 mM HEPES pH 7.5, 75 mM NaCl, 1 mM DTT, 10% glycerol) and heavy buffer (20 mM HEPES pH 7.5, 75 mM NaCl, 1 mM DTT, 40% glycerol, 0.15% glutaraldehyde) were used to generate a 10-40% glycerol gradient with a 0-0.15% glutaraldehyde gradient. Fractions from the gradient were checked via native PAGE. Fractions containing the SIRT6-nucleosome complex were concentrated and buffer exchanged into EM buffer (12.5 mM HEPES pH 7.5, 60 mM KCl, 1.5% glycerol, 1 mM DTT for SIRT6-172 base pair nucleosome and 20 mM HEPES pH 7.5, 75 mM NaCl, 1 mM DTT for SIRT6-145 base pair nucleosome). The final concentration of the SIRT6-nucleosome samples was ~3 μM.

Cryo-EM grids of the complexes were prepared using an established procedure (*38*). Specifically, 3.5 μL of the concentrated sample was applied to copper Quantifoil 2/2 Cu200 (172 base pair sample) or 1.2/1.3 Cu300 (145/147 base pair samples) mesh grids in a FEI Vitrobot Mark IV maintained at 4°C with 100% humidity. The sample was blotted for 3.5 s with a blot force of −1 and then plunge-frozen into liquid ethane.

### Cryo-EM data collection

The SIRT6-147 base pair nucleosome dataset was collected at the Penn State Cryo-Electron Microscopy Facility on a Titan Krios operated at 300 keV and equipped with an FEI Falcon 3 direct electron detector operated in counting mode. 524 movies were collected at 59,000x magnification, which corresponds to 1.14 Å/pixel, at a defocus range of −0.5 to −2.5 μm with ~58 e/Å^2^ accumulated exposure divided between 44 frames **(Table S1)**.

The SIRT6-145 base pair nucleosome dataset was collected at the Pacific Northwest Cryo-EM Center on a Titan Krios operated at 300 keV and equipped with a Gatan K3 direct electron detector and an energy filter (20 eV slit). 7,730 movies were collected in counting mode at 81,000x magnification, which corresponds to 1.059 Å/pixel, at a defocus range of −0.8 to −2.2 μm with ~50 e/Å^2^ accumulated exposure divided between 44 frames **(Table S1)**.

The SIRT6-172 base pair nucleosome dataset was collected at the National Cancer Institute using a Titan Krios operated at 300 keV and equipped with a Gatan K3 direct electron detector and an energy filter (20 eV slit). The data was collected over the course of two separate sessions at 81,000x magnification, which corresponds to 1.08 Å/pixel (0.54 Å/pixel in Super-Resolution mode), at a defocus range of −1.0 to −2.2 μm with ~50 e/Å^2^ accumulated exposure fractionated into 40 super-resolution frames. In total, 11,872 movies were collected over the two sessions (5,122 movies in the first session and 6,750 movies in the second session; **Table S1**).

### Cryo-EM data processing

The SIRT6-172 base pair nucleosome dataset was imported into RELION v3.1 (*39, 40*). Using UCSF MotionCor2 v1.4.1 (*41*) within RELION, raw movies were motion-corrected, binned 2x to 1.08 Å/pixel, and the resulting dose-weighted micrographs (*42*) were imported into cryoSPARC (*43*). *Patch CTF estimation (multi*) was used to estimate defocus values. A nucleosome map was used to generate templates for template picking of particles; these particles were then extracted using a 300 pixel box Fourier-binned to 100 pixels (resulting in 3.24 Å/pixel). 2D classification was performed and classes containing obvious junk were removed. Ab-initio reconstruction was performed with the remaining particles to generate input classes for heterogeneous refinement. Subsequently, a cleaned dataset of nucleosomal the particles was subjected to numerous rounds of 3D classification, yielding a single class with a strong density for SIRT6. To further the quality of the complex, 3D classification without alignment in RELION was performed with a mask focused on the globular domain of SIRT6 and the adjacent DNA. A class with 71,603 particles that represented the most stable positioning of SIRT6 on the nucleosome visually was selected. These particles were refined using non-uniform refinement (*44*) in cryoSPARC, as well as with RELION and with cisTEM (*45*). All reconstructions looked reasonable, but the cisTEM reconstruction was used for further interpretation.

Subsequently, Bayesian polishing was performed in RELION to further improve the quality of the map (*46*). Two rounds of Bayesian polishing and CTF refinement were performed using 1,007,638 particles from an earlier 2.83 Å consensus reconstruction. This improved the resolution of the consensus reconstruction to 2.63 Å. Using these optimized particles, the 71,603-particle subset was refined to 3.07 Å with improved quality of SIRT6 **(Figs. S1 & S2; Table S1)**.

The 147 base pair dataset containing 524 movies was first motion-corrected using cryoSPARC’s *Patch motion correction* (*multi*). Defocus values were calculated using *Patch CTF estimation* (*multi*), and 398,186 particles were blob picked using a sphere with a radius of 120-140 Å and extracted in a 256 pixel box. Five models were generated using ab initio reconstruction and the particles were subjected to heterogeneous refinement. One class was selected based on the presence of SIRT6 and refined to 4.84 Å using non-uniform refinement. These particles were then subjected to classification with the best reconstruction from the previous step seeded three times. One of the resulting maps exhibited a more stable SIRT6-nucleosome complex and was subjected to non-uniform refinement that yielded a reconstruction at 5.0 Å. Using these particles, we re extracted the subset in a larger 300 pixel box; this yielded 40,834 particles that we refined the 4.9 Å final resolution using *Non-Uniform Refinement (New*) **(Fig. S4; Table S1)**.

The SIRT6-145 bp nucleosome dataset was processed as described for SIRT6-147 bp nucleosome datasets, except for motion correction – here, we used UCSF MotionCor2 instead. We obtained two final reconstructions, differing in their SIRT6-interacting DNA: a) 3.28 Å reconstruction from 31,802 particles **(Fig S5C-E, I; Table S1)** and b) 3.27Å reconstruction from 34,737 particles **(Fig S5F-H,J; Table S1)**.

The final resolutions reported were calculated using Fourier Shell Correlation (FSC) and assessed at the 0.143 cutoff following gold-standard refinement (*47*). All file format conversions between cryoSPARC and RELION, as well as particle stack preparation for cisTEM refinement were performed using UCSF pyem v0.5 (*48*).

### Model building and refinement

PDB 3LZ0 (*49*) was used as a starting model for the nucleosome, and 3PKI (*13*) and 7CL0 (*19*) were used as starting models for SIRT6. The models were rigid-body fitted into the 3.07 Å reconstruction using the “fit in map” function in UCSF ChimeraX (*50*) and then further optimized in Coot (*51*).

A combination of 3D Coulomb potential maps from the 172 base pair dataset were used to construct the final model. Using the highest resolved 2.63 Å reconstruction in Coot, histones and core DNA were optimized and the SIRT6 acidic patch-interacting loop was constructed. The 3.07 Å reconstruction was then used to position and adjust the SIRT6 globular domain, zinc-binding domain, histone H2A C-terminal tail, histone H3 N-terminal tail, and the DNA overhang. The final model was refined using phenix.real_space_refine (*52*) with secondary structure, Ramachandran, and rotamer restraints. Lastly, the model was validated manually in Coot and with Molprobity (*53*) using comprehensive validation (cryo-EM) in Phenix. Model statistics are reported in **Table S2**.

### Electrophoresis Mobility Shift Assays (Nucleosome Binding Assays)

10 μL reactions of SIRT6 variants at indicated concentrations were incubated with 100 nM of wild-type NCPs for 30 min at 30°C in 5x binding buffer (25 mM HEPES pH 7.3, 150 mM NaCl, 4.5 mM MgCl_2_, 1 mM DTT, 0.01% Tween, 0.1 mg/ml BSA). Reactions were quenched with 2 μL of 50% glycerol and electrophoresed on 4.5% native PAGE gels at 100 V, 4 °C.

### Histone Deacetylation Assays

Wild-type SIRT6 (500nM or 125nM) and SIRT6(R175A) (200nM) was incubated with nucleosomes (125nM) containing either acetylated H3K9, H3K56, or H3K27, and/or with truncated H2A (H2At) at 37 °C for varying time points in HDAC reaction buffer (25mM HEPES pH 7.3, 49 mM NaCl, 4.5mM MgCl_2_, 1mM DTT). Reactions were quenched with the addition of 2X SDS sample buffer, boiled, and then electrophoresed on a 12 % SDS-PAGE gel. Proteins on the gel were transferred to a PVDF membrane (Millipore IPFL 00010), which was first blocked with 10 % milk, and then probed with either 1:10,000 anti-H3K9ac antibody (Active Motif #39038), 1:1,000 anti-H3K56ac antibody (Active Motif #39133), or 1:3,000 anti-H3K27ac antibody (Active Motif #39082), and also probed with 1:3,000 anti-H2B antibody (Abcam #64039). Western blots were quantified using ImageJ (*54*) or Bio-Rad Image Lab.

## Supporting information

Supplementary File on its own

## Acknowledgments

We thank Szu-Yu Kuan and Eric Baron at Penn State for nucleosome preparations, Adam Wier at NCI, Theo Humphreys at PNCC for assistance with EM data collection, Joseph Cho and Carol Bator of the Penn State Cryo-Electron Microscopy Facility for assistance with EM screening and data collection, and Jennifer Gray at Penn State for assistance with training on the T20 for negative stain EM. This work was supported by:

National Institutes of Health grant F32GM137463 (USC)

National Institutes of Health grant T32BM107000 (ARB)

National Institutes of Health grant R35GM122519 (CLP)

National Institutes of Health grant R35GM127034 (ST)

This project is funded, in part, under a grant from the Pennsylvania Department of Health using Tobacco CURE Funds. The Department specifically disclaims responsibility for any analyses, interpretations or conclusion.

The figures for this manuscript were generated using PyMOL and UCSF ChimeraX. UCSF ChimeraX is developed by the Resource for Biocomputing, Visualization, and Informatics at the University of California, San Francisco, with support from National Institutes of Health R01-GM129325 and the Office of Cyber Infrastructure and Computational Biology, National Institute of Allergy and Infectious Diseases.

## Author contributions

Conceptualization: CLP, ST, JPA

Methodology: USC, OR, AB, JZ, JLF, CLP, ST, JPA

Investigation: USC, OR, AB, JZ, JLF, JPA

Visualization: ST, JPA

Funding acquisition: USC, CLP, ST, JPA

Project administration: CLP, ST, JPA

Supervision: CLP, ST, JPA

Writing – original draft: USC, ST, JPA

Writing – review & editing: OR, AB, CLP, ST, JPA

## Competing interests

Authors declare that they have no competing interests.

## Data and materials availability

The atomic coordinates of the SIRT6-nucleosome complex have been deposited to the RCSB Protein Data Bank with PDB ID XXXX. The cryo-EM Coulomb potential maps were deposited in the Electron Microscopy Data Bank as EMD-XXXXX (best-resolved SIRT6-nucleosome complex), EMD-XXXXY (SIRT6 in position 2), EMD-XXXYX (SIRT6 in position 3). The raw cryo-EM data was deposited in EMPIAR (EMPIAR-ZZZ).

## Supplementary Materials

**Fig. S1.**
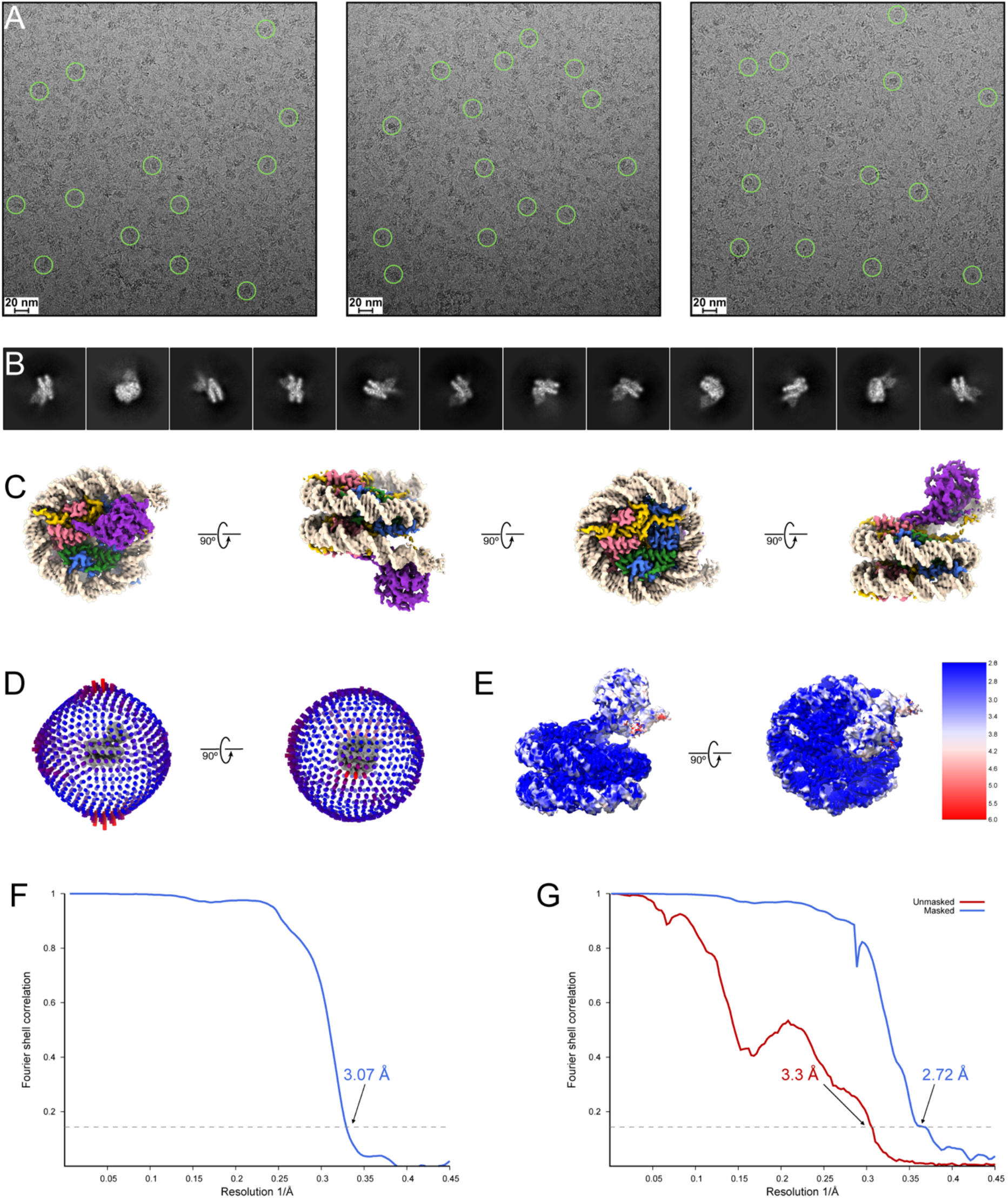
Cryo-EM studies of SIRT6 bound to 172 bp nucleosome. **(A)** Representative motion-corrected micrographs from the SIRT6 bound to 172 bp nucleosome dataset. **(B)** Representative 2D classes of SIRT6 bound to 172 bp nucleosome. **(C)** Four different views of the cryo-EM map of SIRT6 bound to 172 bp nucleosome generated with cisTEM. The structure is color-coded with histone H3 in light blue, histone H4 in light green, histone H2A in light yellow, histone H2B in light pink, DNA strands in light/dark gray, and SIRT6 in dark blue. **(D)** Angular distribution of particles used to generate the cryo-EM map of SIRT6 bound to 172 bp nucleosome in cisTEM. **(E)** Cryo-EM map of SIRT6 bound to 172 bp nucleosome generated in cisTEM colored by estimated local resolution determined with FSC = 0.143 cutoff in cryoSPARC. **(F)** Fourier shell correlation curve between two half-maps for the SIRT6 bound to 172 bp nucleosome refinement determined by cisTEM. **(G)** Unmasked (red) and masked (blue) Fourier shell correlation curves between two independent half-maps for the SIRT6 bound to 172 bp nucleosome refinement determined by cryoSPARC.

**Fig. S2:**
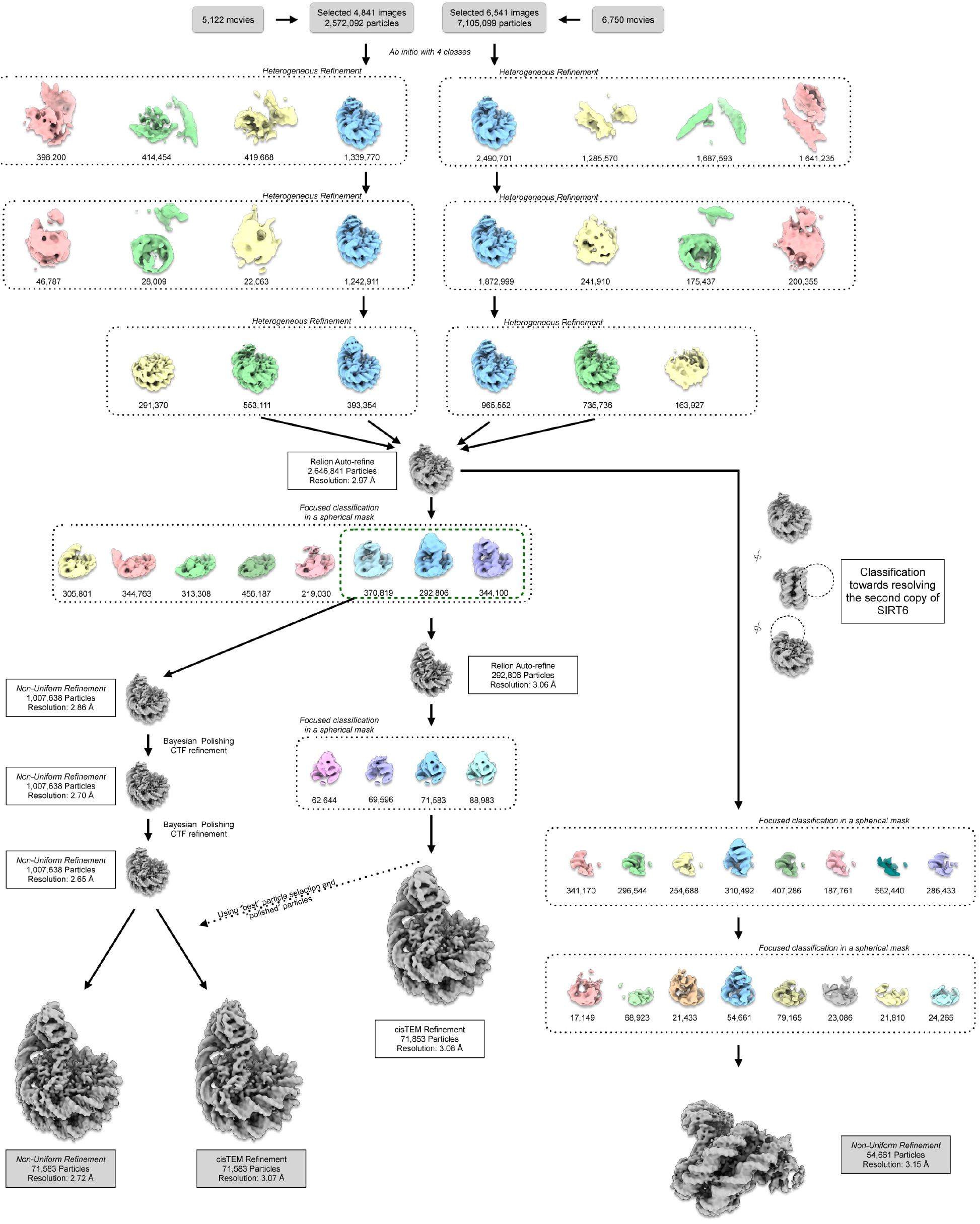
Data processing for SIRT6 bound to 172 bp nucleosome. Flowchart for cryo-EM data processing of the SIRT6 bound to 172 bp nucleosome datasets as described in Methods. Initial processing steps were performed in cryoSPARC with the number of particles moving into each step noted. Focused classifications without alignment were performed in RELION-3.1 to improve SIRT6 density. Rounds of Bayesian polishing and CTF refinement were also performed in RELION-3.1 to improve the overall map. Final refinements were performed in both cryoSPARC and cisTEM.

**Fig. S3:**
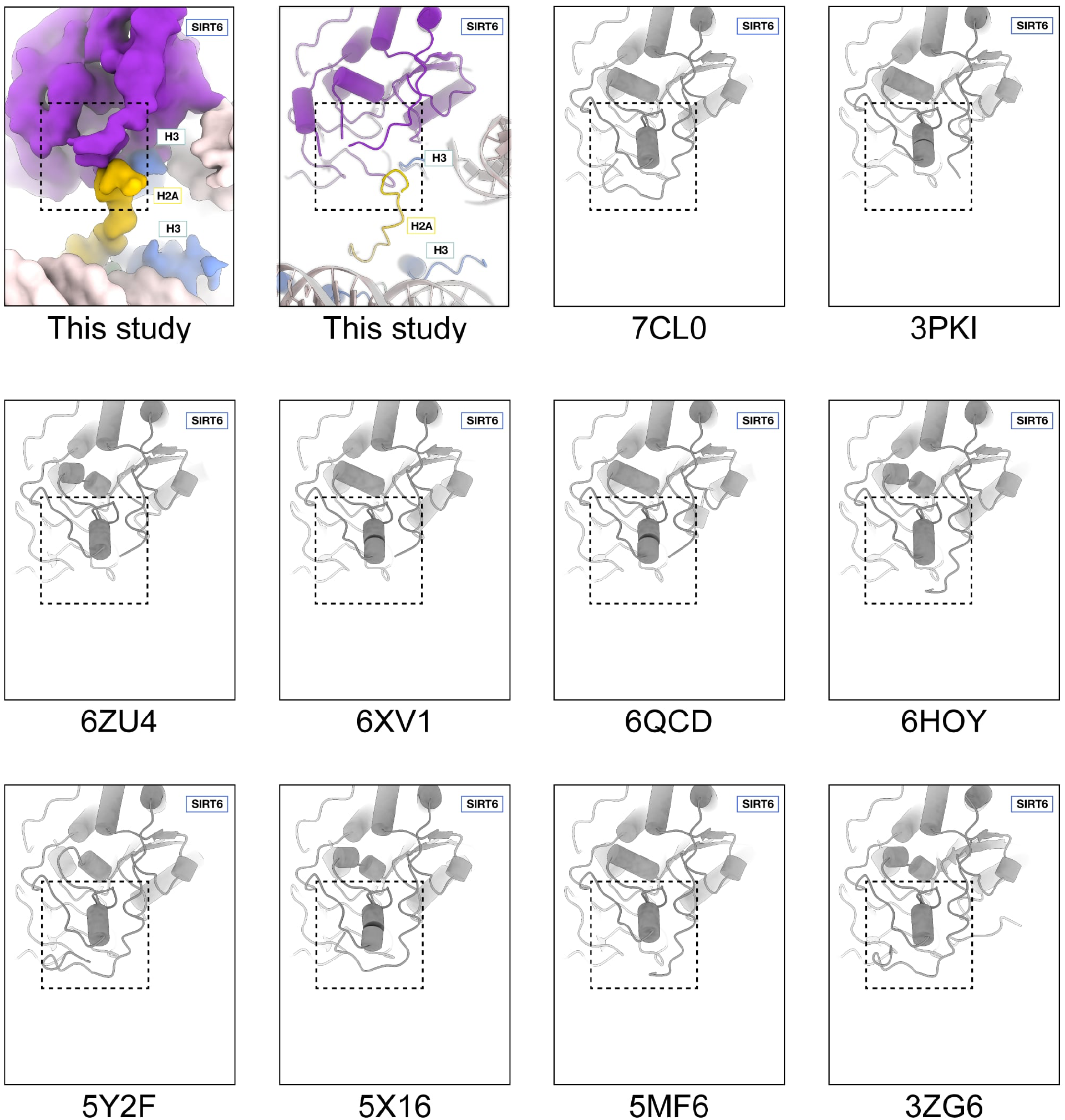
Comparison of SIRT6 residues 64-80 between structures. SIRT6 is missing density for residues 64-80 in our structure (“this study” panels) suggesting the region is disordered. SIRT6 crystal structures have their PDBID codes indicated underneath. Residues 64-80 are ordered and form a small alpha-helix in all crystal structures of SIRT6 with ADP ribose.

**Fig. S4:**
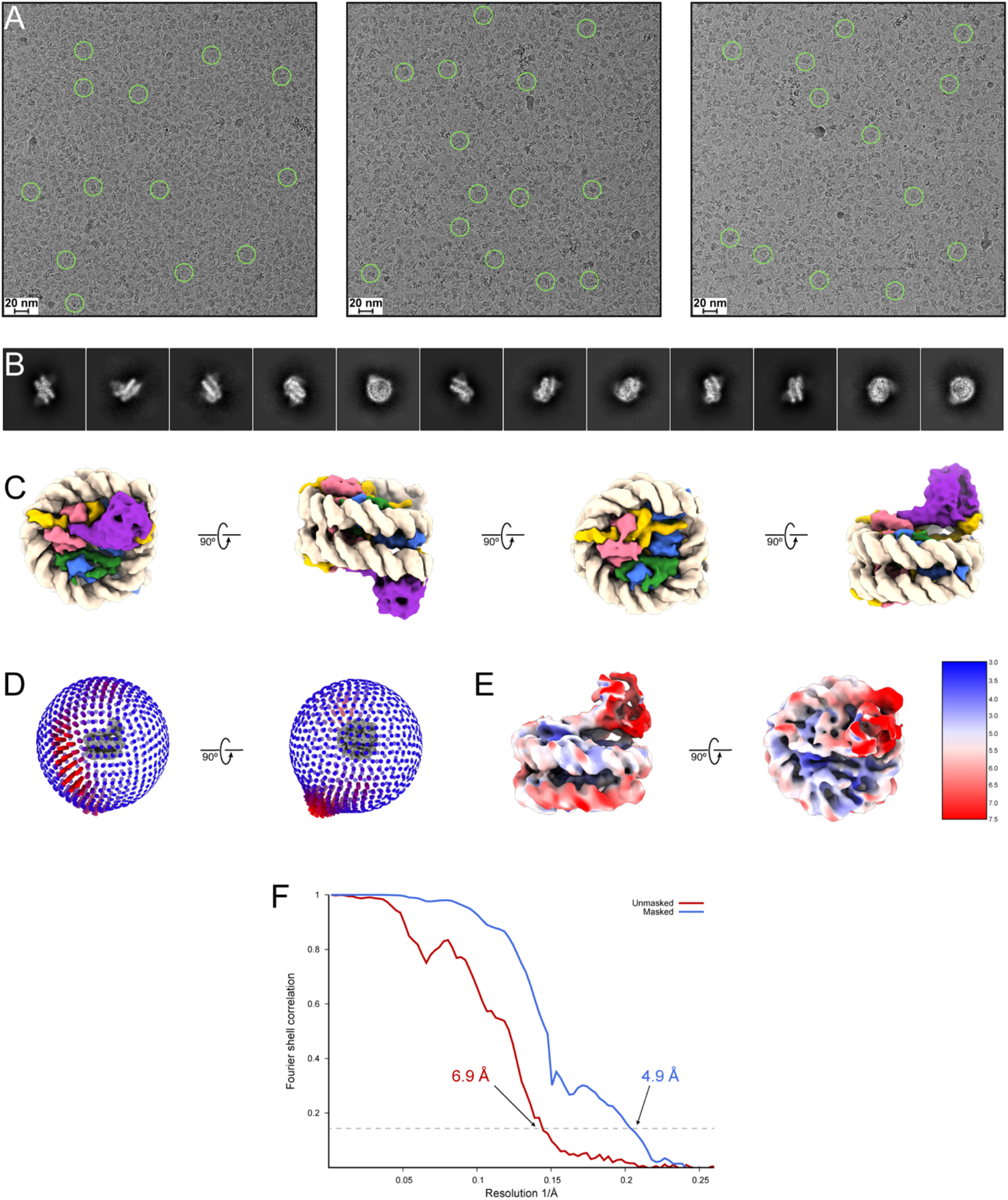
Cryo-EM studies of SIRT6 bound to 147 bp nucleosome. **(A)** Representative motion-corrected micrographs from the SIRT6 bound to 147 bp nucleosome dataset. **(B)** Representative 2D classes of SIRT6 bound to 147 bp nucleosome. **(C)** Four different views of the cryo-EM map of SIRT6 bound to 147 bp nucleosome. The structure is color-coded with histone H3 in light blue, histone H4 in light green, histone H2A in light yellow, histone H2B in light pink, DNA strands in light/dark gray, and SIRT6 in dark blue. **(D)** Angular distribution of particles used to generate the cryo-EM map of SIRT6 bound to 147-base pair nucleosome. **(E)** Cryo-EM map of SIRT6 bound to 147 bp nucleosome colored by estimated local resolution determined with FSC = 0.143 cutoff in cryoSPARC. **(F)** Unmasked (red) and masked (blue) Fourier shell correlation curves between two independent half-maps for the SIRT6 bound to 147 bp nucleosome refinement determined by cryoSPARC.

**Fig. S5:**
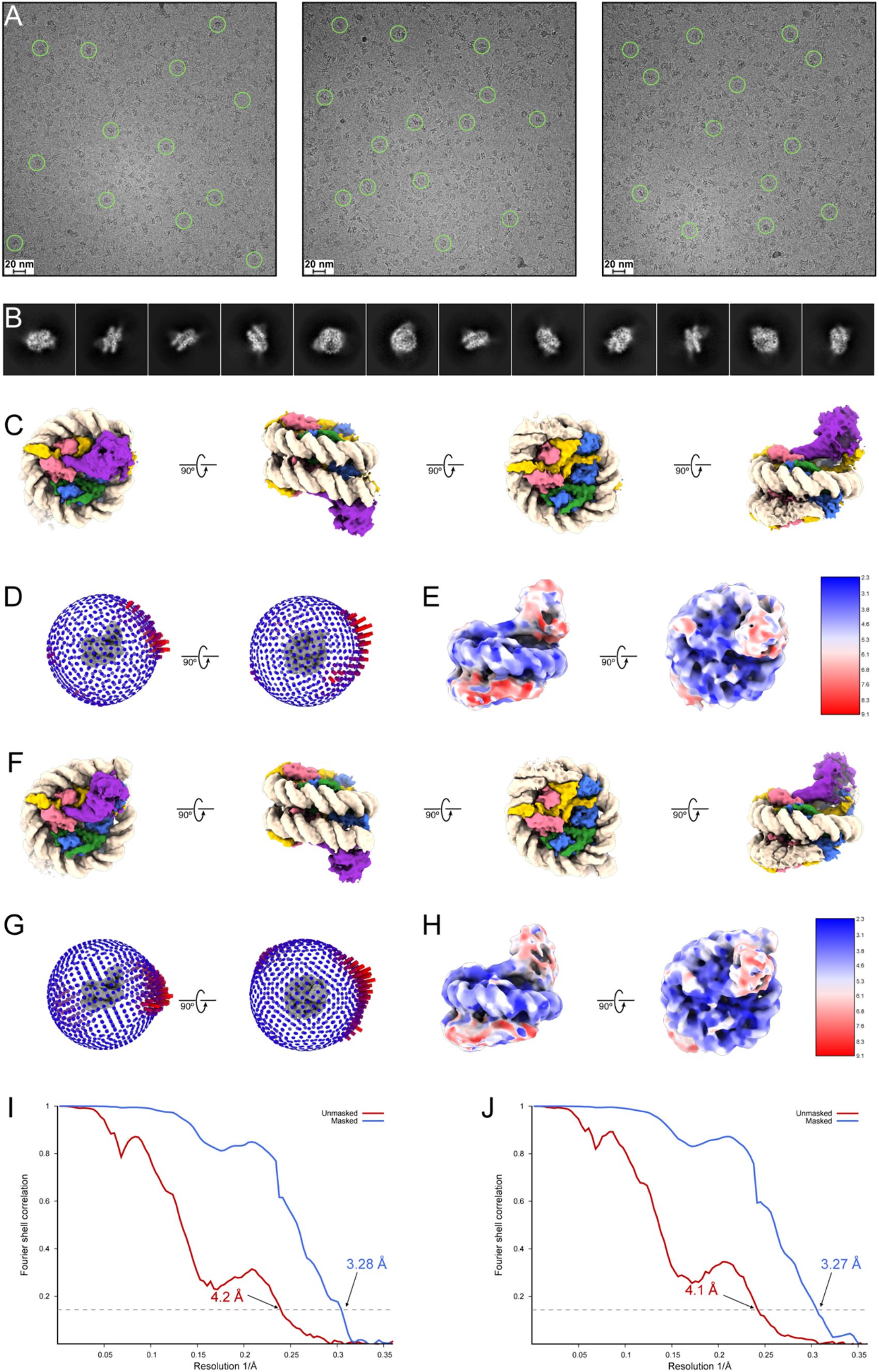
Cryo-EM studies of SIRT6 bound to 145 bp nucleosome. **(A)** Representative motion-corrected micrographs from the SIRT6 bound to 145 bp nucleosome dataset. **(B)** Representative 2D classes of SIRT6 bound to 145 bp nucleosome. **(C)** Four different views of one cryo-EM map determined of SIRT6 bound to 145 bp nucleosome. The structure is color-coded with histone H3 in light blue, histone H4 in light green, histone H2A in light yellow, histone H2B in light pink, DNA strands in light/dark gray, and SIRT6 in dark blue. **(D)** Angular distribution of particles used to generate the first cryo-EM map of SIRT6 bound to 145 bp nucleosome. **(E)** The first cryo-EM map of SIRT6 bound to 145 bp nucleosome colored by estimated local resolution determined with FSC = 0.143 cutoff in cryoSPARC. **(F)** Four different views of the second cryo-EM map determined of SIRT6 bound to 145 bp nucleosome. The structure is color-coded as in (C). **(G)** Angular distribution of particles used to generate the second cryo-EM map of SIRT6 bound to 145 bp nucleosome. **(H)** The second cryo-EM map of SIRT6 bound to 145 bp nucleosome colored by estimated local resolution determined with FSC = 0.143 cutoff in cryoSPARC. **(I)** Unmasked (red) and masked (blue) Fourier shell correlation curves between two independent half-maps for the first SIRT6 bound to 145 bp nucleosome refinement determined by cryoSPARC. **(J)** Unmasked (red) and masked (blue) Fourier shell correlation curves between two independent half-maps for the second SIRT6 bound to 145 bp nucleosome refinement determined by cryoSPARC.

**Table S1.**
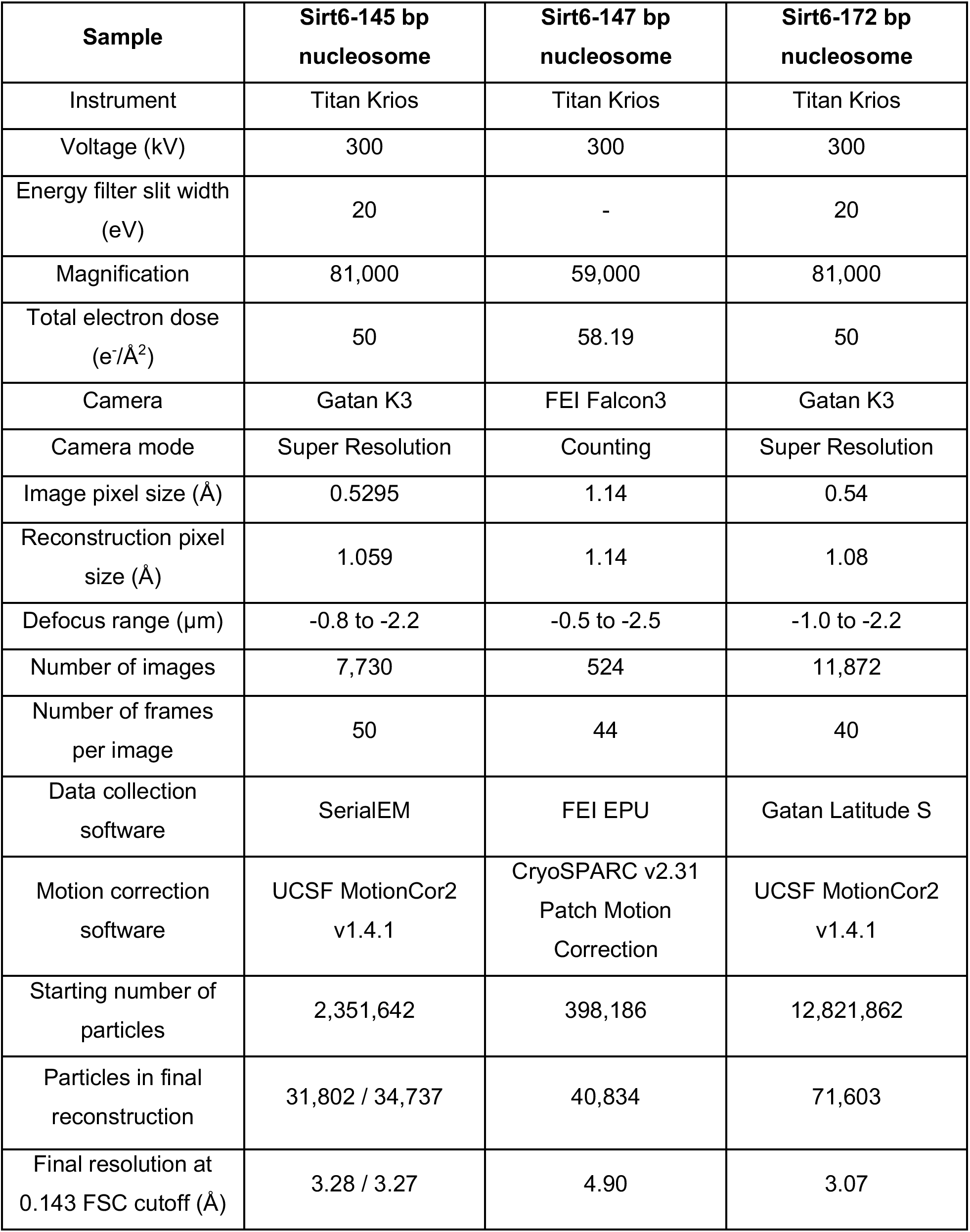
Summary for cryo-EM data collection and refinement.

**Table S2.**
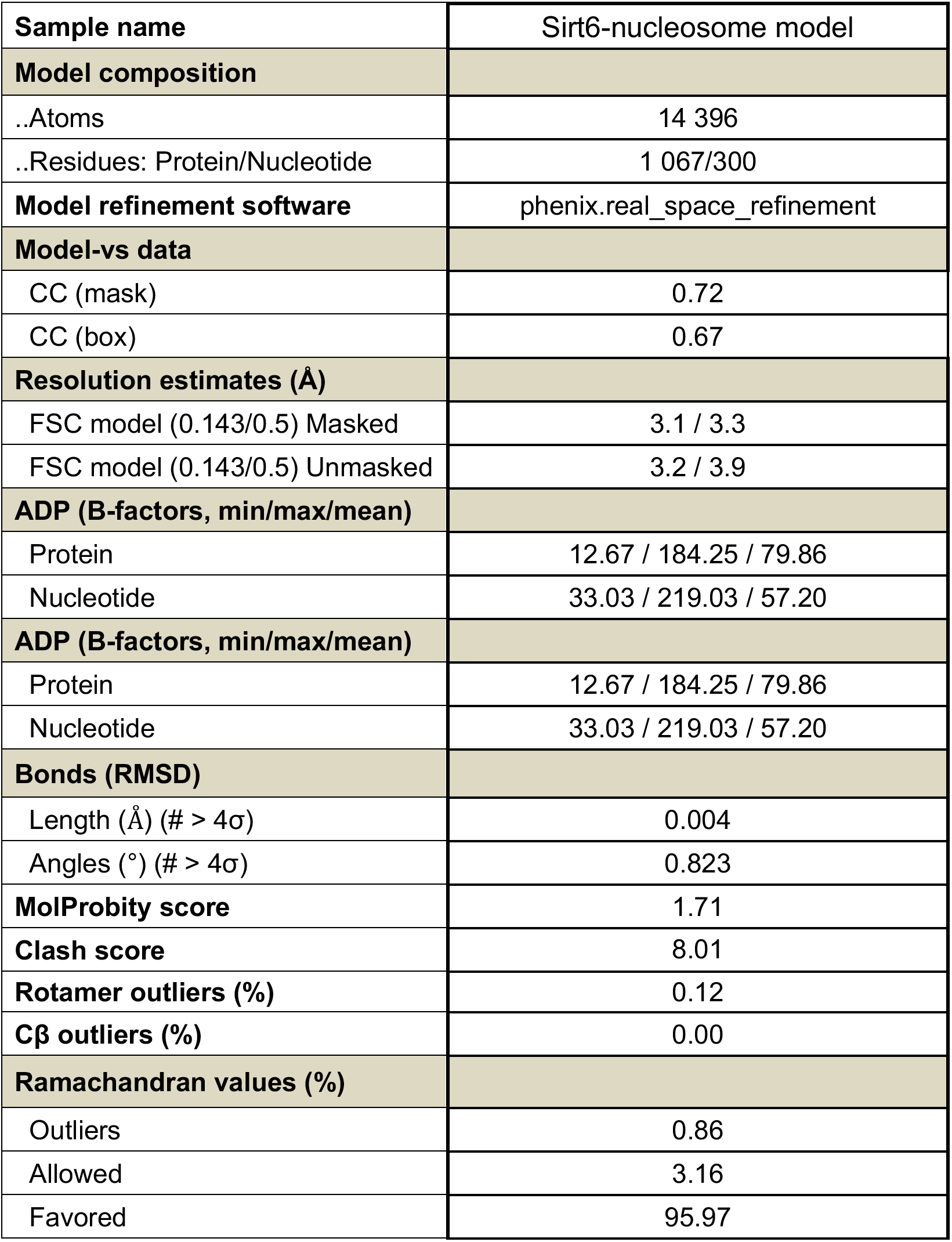
Model refinement statistics.

## Notes

### Competing Interest Statement

The authors have declared no competing interest.

