## Supplementary File on its own for "Cryo-EM structure of the human Sirtuin 6-nucleosome complex"

**This PDF file includes:**

Figs. S1 to S5
Tables S1 to S2

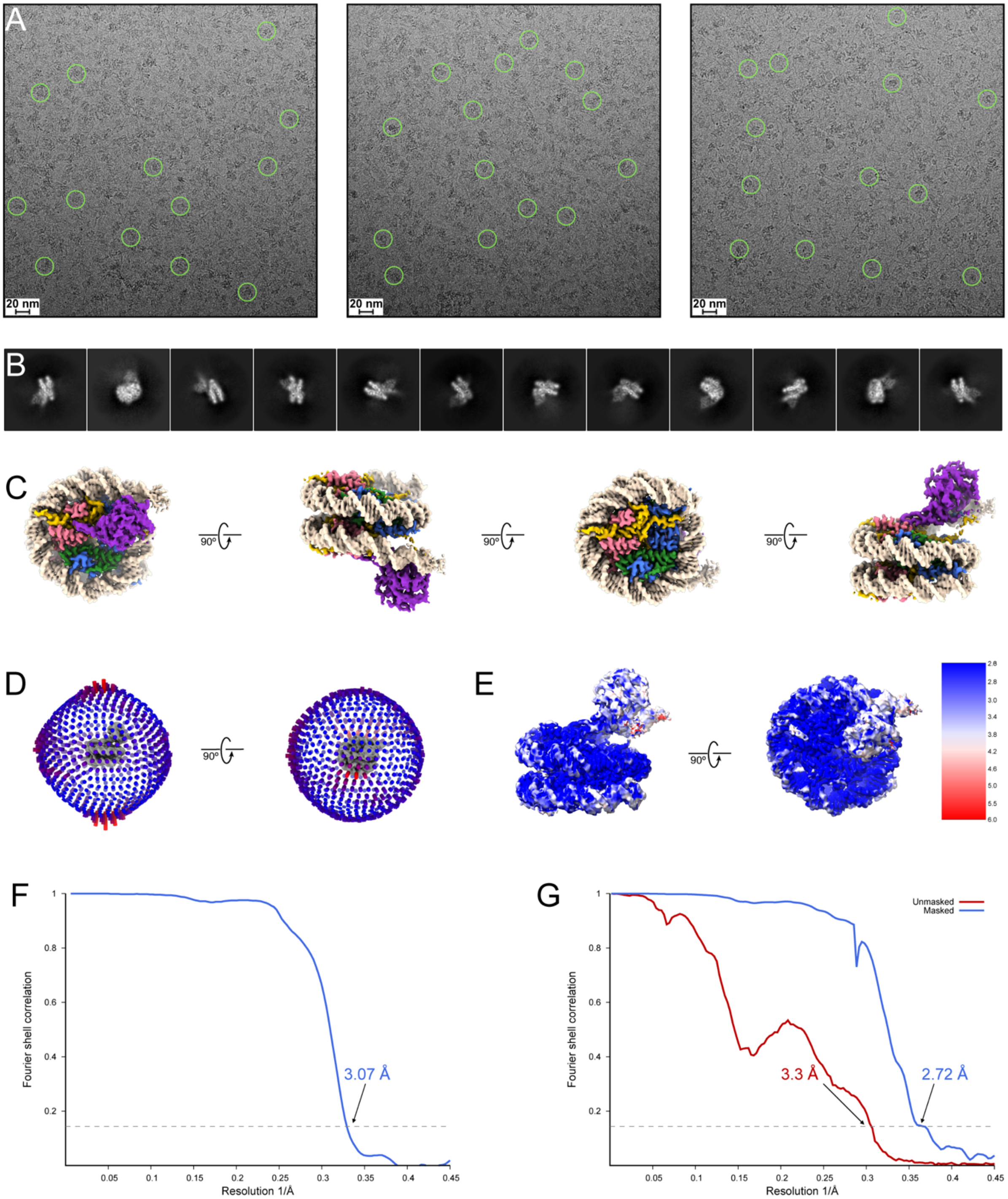


Fig. S1. Cryo-EM studies of SIRT6 bound to 172 bp nucleosome. **(A)** Representative motion-corrected micrographs from the SIRT6 bound to 172 bp nucleosome dataset. **(B)** Representative 2D classes of SIRT6 bound to 172 bp nucleosome. **(C)** Four different views of the cryo-EM map of SIRT6 bound to 172 bp nucleosome generated with cisTEM. The structure is color-coded with histone H3 in light blue, histone H4 in light green, histone H2A in light yellow, histone H2B in light pink, DNA strands in light/dark gray, and SIRT6 in dark blue. **(D)** Angular distribution of particles used to generate the cryo-EM map of SIRT6 bound to 172 bp nucleosome in cisTEM.

**(E)** Cryo-EM map of SIRT6 bound to 172 bp nucleosome generated in cisTEM colored by estimated local resolution determined with FSC = 0.143 cutoff in cryoSPARC. **(F)** Fourier shell correlation curve between two half-maps for the SIRT6 bound to 172 bp nucleosome refinement determined by cisTEM. **(G)** Unmasked (red) and masked (blue) Fourier shell correlation curves between two independent half-maps for the SIRT6 bound to 172 bp nucleosome refinement determined by cryoSPARC.

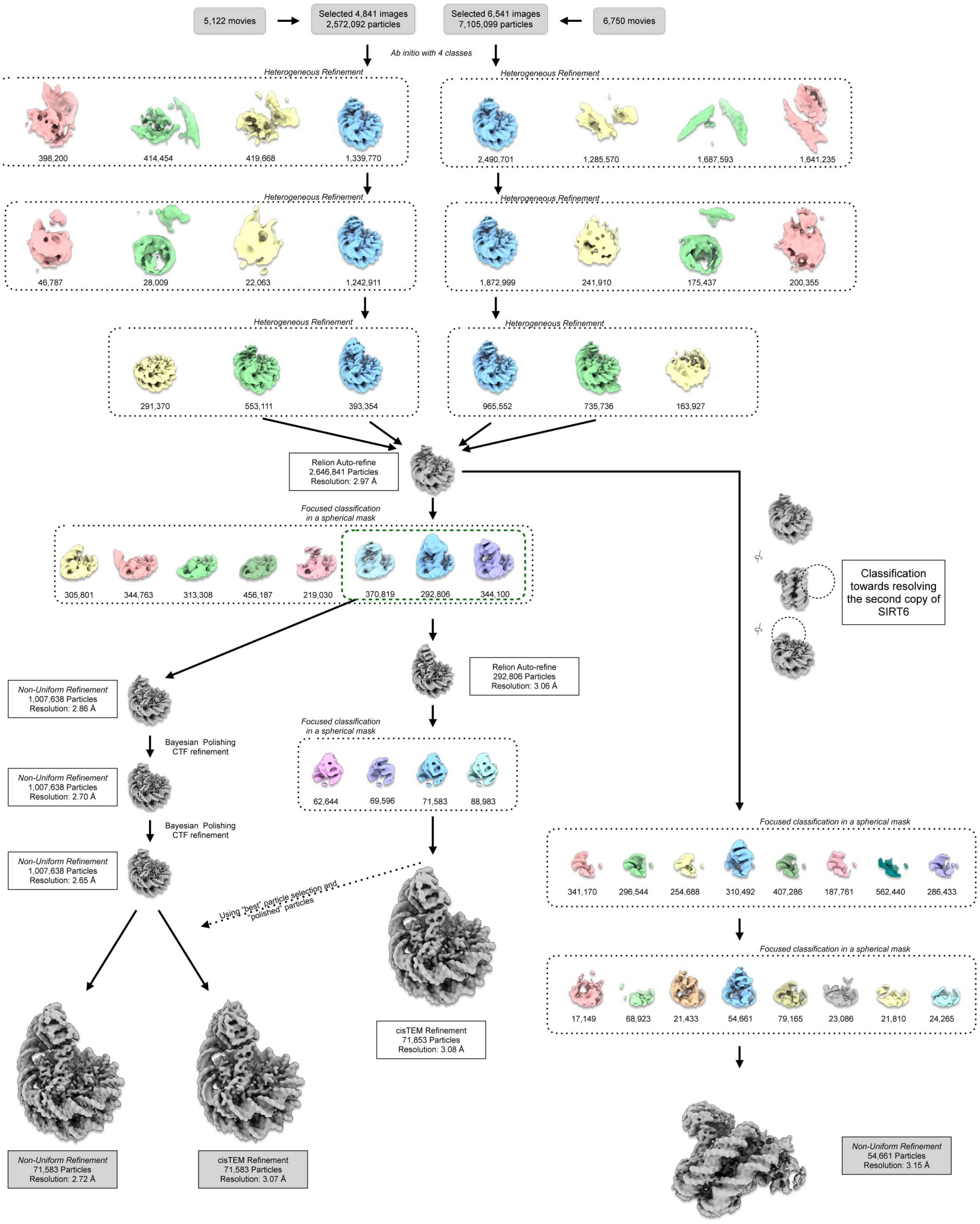


Fig. S2: Data processing for SIRT6 bound to 172 bp nucleosome. Flowchart for cryo-EM data processing of the SIRT6 bound to 172 bp nucleosome datasets as described in Methods. Initial processing steps were performed in cryoSPARC with the number of particles moving into each step noted. Focused classifications without alignment were performed in RELION-3.1 to improve SIRT6 density. Rounds of Bayesian polishing and CTF refinement were also performed in RELION-3.1 to improve the overall map. Final refinements were performed in both cryoSPARC and cisTEM.

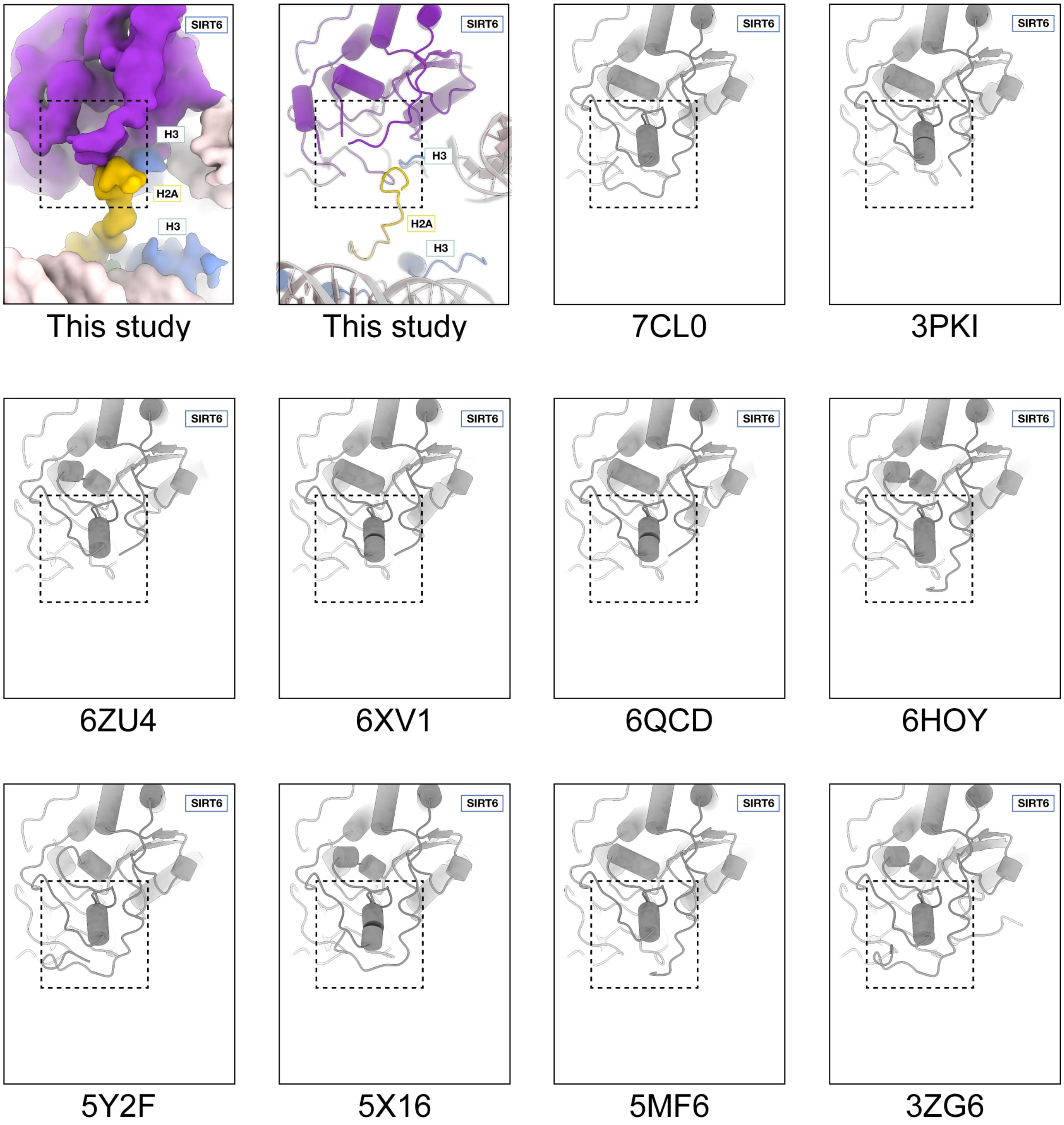


Fig. S3: Comparison of SIRT6 residues 64-80 between structures. SIRT6 is missing density for residues 64-80 in our structure ("this study" panels) suggesting the region is disordered. SIRT6 crystal structures have their PDBID codes indicated underneath. Residues 64-80 are ordered and form a small alpha-helix in all crystal structures of SIRT6 with ADP ribose.

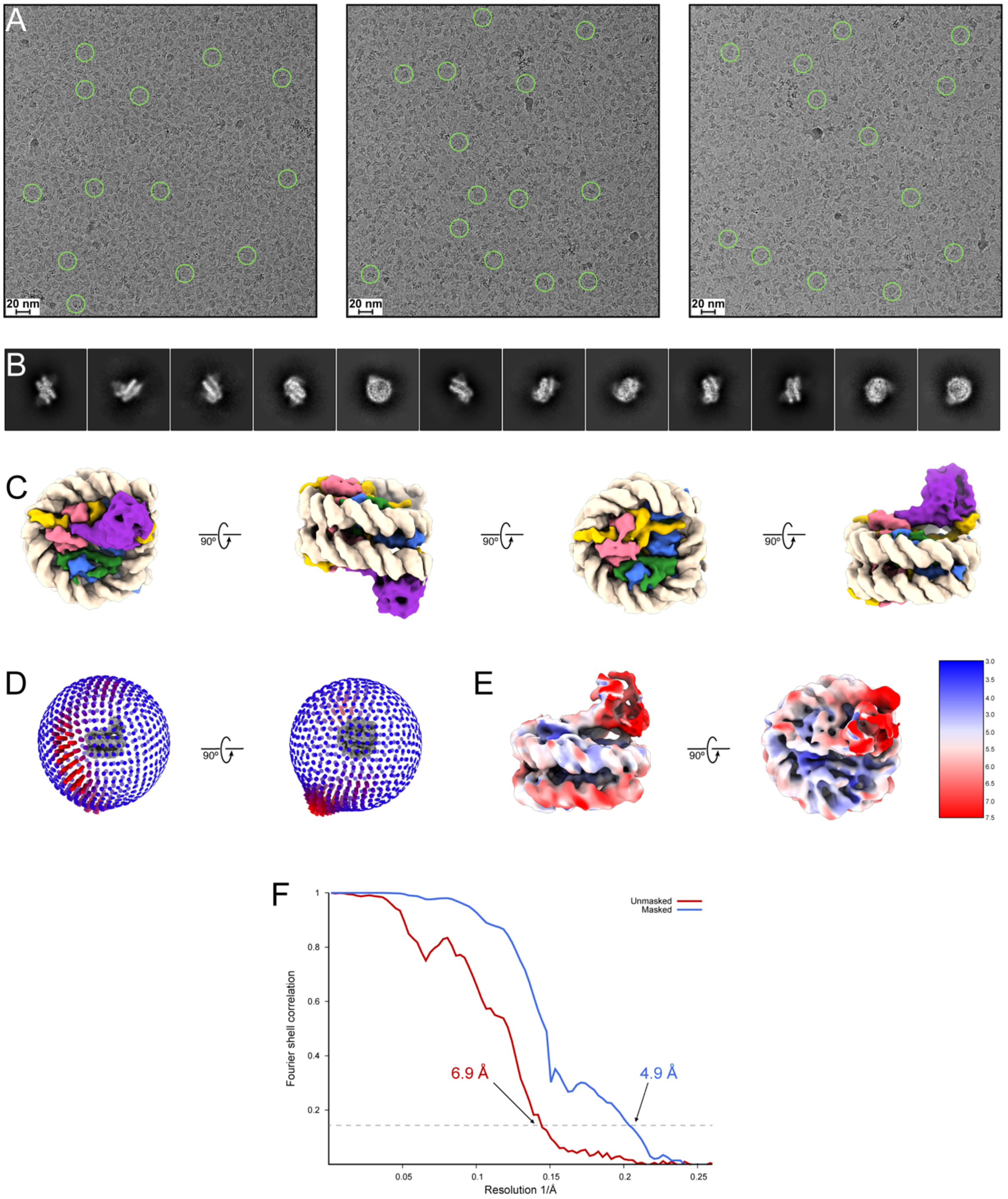


Fig. S4: Cryo-EM studies of SIRT6 bound to 147 bp nucleosome. **(A)** Representative motion-corrected micrographs from the SIRT6 bound to 147 bp nucleosome dataset. **(B)** Representative 2D classes of SIRT6 bound to 147 bp nucleosome. **(C)** Four different views of the cryo-EM map of SIRT6 bound to 147 bp nucleosome. The structure is color-coded with histone H3 in light blue, histone H4 in light green, histone H2A in light yellow, histone H2B in light pink, DNA strands in light/dark gray, and SIRT6 in dark blue. **(D)** Angular distribution of particles used to generate the cryo-EM map of SIRT6 bound to 147-base pair nucleosome. **(E)** Cryo-EM map of SIRT6 bound to 147 bp nucleosome colored by estimated local resolution determined with FSC = 0.143 cutoff in cryoSPARC. **(F)** Unmasked (red) and masked (blue) Fourier shell correlation curves between two independent half-maps for the SIRT6 bound to 147 bp nucleosome refinement determined by cryoSPARC.

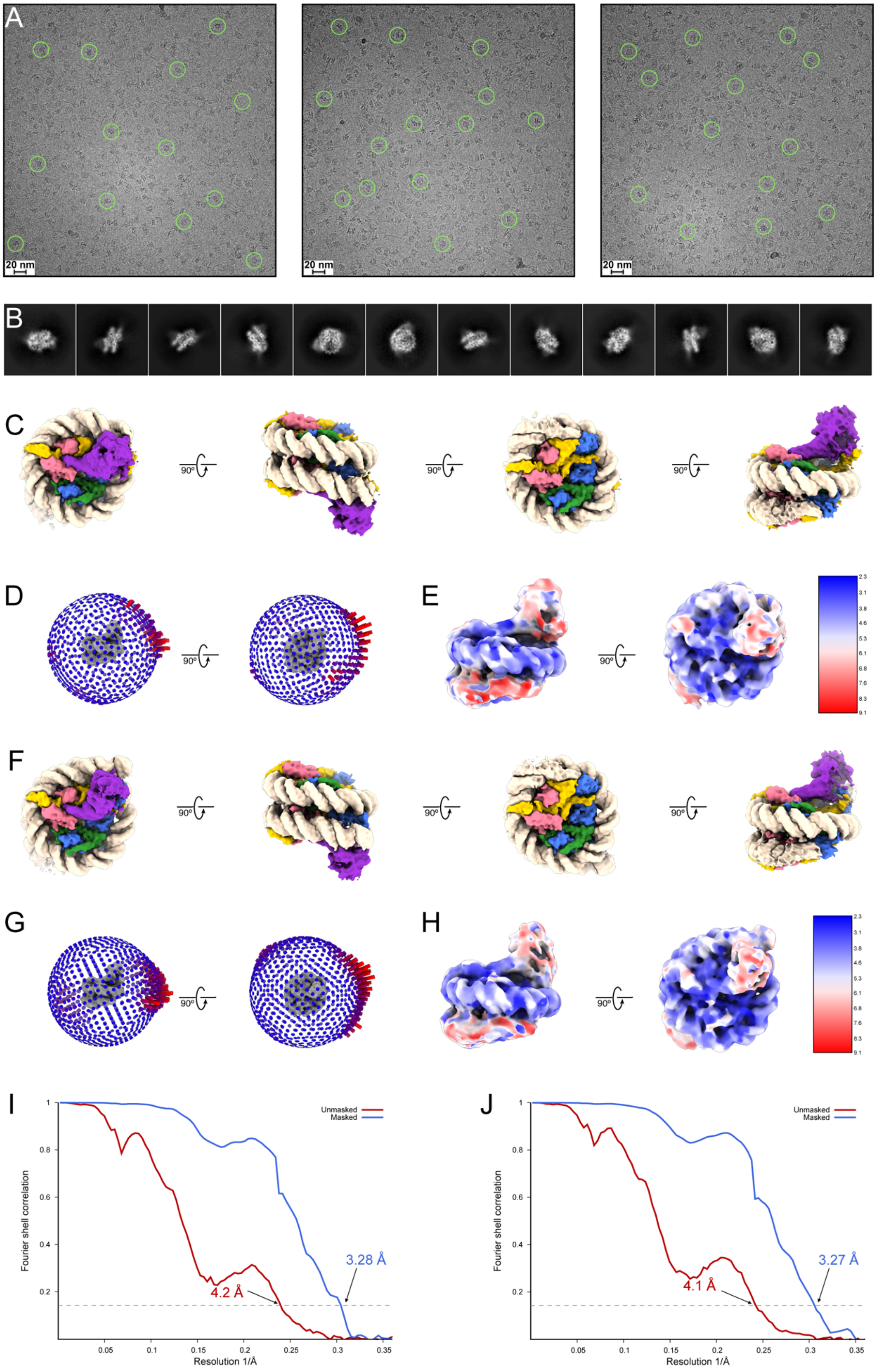
A
20 nm
20 nm
20 nm
B
C
90°
90°
90°
D
90°
E
90°
F
90°
90°
90°
G
90°
H
90°
I
Unmasked
Masked
Fourier shell correlation
Resolution 1/Å
3.28 Å
4.2 Å
J
Unmasked
Masked
Fourier shell correlation
Resolution 1/Å
3.27 Å
4.1 Å

Fig. S5: Cryo-EM studies of SIRT6 bound to 145 bp nucleosome. **(A)** Representative motion-corrected micrographs from the SIRT6 bound to 145 bp nucleosome dataset. **(B)** Representative 2D classes of SIRT6 bound to 145 bp nucleosome. **(C)** Four different views of one cryo-EM map determined of SIRT6 bound to 145 bp nucleosome. The structure is color-coded with histone H3 in light blue, histone H4 in light green, histone H2A in light yellow, histone H2B in light pink, DNA strands in light/dark gray, and SIRT6 in dark blue. **(D)** Angular distribution of particles used to generate the first cryo-EM map of SIRT6 bound to 145 bp nucleosome. **(E)** The first cryo-EM map of SIRT6 bound to 145 bp nucleosome colored by estimated local resolution determined with FSC = 0.143 cutoff in cryoSPARC. **(F)** Four different views of the second cryo-EM map determined of SIRT6 bound to 145 bp nucleosome. The structure is color-coded as in (C). **(G)** Angular distribution of particles used to generate the second cryo-EM map of SIRT6 bound to 145 bp nucleosome. **(H)** The second cryo-EM map of SIRT6 bound to 145 bp nucleosome colored by estimated local resolution determined with FSC = 0.143 cutoff in cryoSPARC. **(I)** Unmasked (red) and masked (blue) Fourier shell correlation curves between two independent half-maps for the first SIRT6 bound to 145 bp nucleosome refinement determined by cryoSPARC. **(J)** Unmasked (red) and masked (blue) Fourier shell correlation curves between two independent half-maps for the second SIRT6 bound to 145 bp nucleosome refinement determined by cryoSPARC.

Table S1. Summary for cryo-EM data collection and refinement

| **Sample** | **Sirt6-145 bp nucleosome** | **Sirt6-147 bp nucleosome** | **Sirt6-172 bp nucleosome** |
|---|---|---|---|
| Instrument | Titan Krios | Titan Krios | Titan Krios |
| Voltage (kV) | 300 | 300 | 300 |
| Energy filter slit width (eV) | 20 | - | 20 |
| Magnification | 81,000 | 59,000 | 81,000 |
| Total electron dose ($e^-/Å^2$) | 50 | 58.19 | 50 |
| Camera | Gatan K3 | FEI Falcon3 | Gatan K3 |
| Camera mode | Super Resolution | Counting | Super Resolution |
| Image pixel size (Å) | 0.5295 | 1.14 | 0.54 |
| Reconstruction pixel size (Å) | 1.059 | 1.14 | 1.08 |
| Defocus range (µm) | -0.8 to -2.2 | -0.5 to -2.5 | -1.0 to -2.2 |
| Number of images | 7,730 | 524 | 11,872 |
| Number of frames per image | 50 | 44 | 40 |
| Data collection software | SerialEM | FEI EPU | Gatan Latitude S |
| Motion correction software | UCSF MotionCor2 v1.4.1 | CryoSPARC v2.31 Patch Motion Correction | UCSF MotionCor2 v1.4.1 |
| Starting number of particles | 2,351,642 | 398,186 | 12,821,862 |
| Particles in final reconstruction | 31,802 / 34,737 | 40,834 | 71,603 |
| Final resolution at 0.143 FSC cutoff (Å) | 3.28 / 3.27 | 4.90 | 3.07 |

Table S2. Model refinement statistics

| **Sample name** | Sirt6-nucleosome model |
|---|---|
| **Model composition** | |
| ..Atoms | 14 396 |
| ..Residues: Protein/Nucleotide | 1 067/300 |
| **Model refinement software** | phenix.real_space_refinement |
| **Model-vs data** | |
| CC (mask) | 0.72 |
| CC (box) | 0.67 |
| **Resolution estimates (Å)** | |
| FSC model (0.143/0.5) Masked | 3.1 / 3.3 |
| FSC model (0.143/0.5) Unmasked | 3.2 / 3.9 |
| **ADP (B-factors, min/max/mean)** | |
| Protein | 12.67 / 184.25 / 79.86 |
| Nucleotide | 33.03 / 219.03 / 57.20 |
| **ADP (B-factors, min/max/mean)** | |
| Protein | 12.67 / 184.25 / 79.86 |
| Nucleotide | 33.03 / 219.03 / 57.20 |
| **Bonds (RMSD)** | |
| Length (Å) (# > 4σ) | 0.004 |
| Angles (°) (# > 4σ) | 0.823 |
| **MolProbity score** | 1.71 |
| **Clash score** | 8.01 |
| **Rotamer outliers (%)** | 0.12 |
| **Cβ outliers (%)** | 0.00 |
| **Ramachandran values (%)** | |
| Outliers | 0.86 |
| Allowed | 3.16 |
| Favored | 95.97 |
